## Supplementary material for "Perception of biological motion in point-light displays by jumping spiders": Full analysis and python script: SI1_full-analysis.html

ESM02 - Data Analysis


### ESM02 - Data Analysis

##### Data Analysis

Abstract

This supplement provides the entire R script and output of the statistical analysis we performed and figures produced, in their original form. It is presented in the spirit of open and transparent science, but has not been carefully curated.

### Setup

#### Load packages

```
library(readODS) #to read raw data
library(glmmTMB) #for mixed models
library(car) #for anova on mixed models
library(DHARMa) #for goodness of fit of the model
library(emmeans) #for post hoc
library(ggplot2) #to plot
```

#### Load data

Will load as separate data frames the main experiment and the control

#### Structure columns

```
main$date <- as.factor(main$date)
main$subj <- as.factor(main$subj)
main$sex <- as.factor(main$sex)
main$trialn <- as.factor(main$trialn)
main$cond <- as.factor(main$cond)
main$corrside <- as.factor(main$corrside)
main$stimpres <- as.factor(main$stimpres)

control$date <- as.factor(control$date)
control$subj <- as.factor(control$subj)
control$sex <- as.factor(control$sex)
control$trialn <- as.factor(control$trialn)
control$cond <- as.factor(control$cond)
control$corrside <- as.factor(control$corrside)
control$stimpres <- as.factor(control$stimpres)
```

| column name | description |
| --- | --- |
| date | date of the experiment |
| subj | subject ID |
| sex | subject sex |
| trialn | trial number, 1 to 4 |
| stimn | stimulus number of the current trial, 0 to 9 |
| cond | condition of the current trial |
| corrside | psition of the expected correct stimulus |
| stimpres | is the stimulus visible on the screen currently |
| framefromonset | frame number from stimulus start |
| timefromonset | time in second from stimulus start |
| stimpos\_rad | position of the stimulus on the screen in radians |
| stimpos\_deg | position of the stimulus on the screen in degrees |
| dirval\_rad | purified Z peak value (with subtracted X and Y), for this frame, expressed in radians/frame. Higher values mean purer Z (less X and Y) and faster rotatons. The peak is positive if coherent with the position of the correct stimulus, negative if towards the wrong |
| absval\_rad | same as dirval\_rad, but all the values are positive, to observe the consistency with stimpos |
| dirval\_deg | same as dirval\_rad, but in degrees/frame |
| absval\_deg | same as absval\_rad, but in degrees/frame |
| dirval\_rad\_s | same as dirval\_rad, but multipied by 120 to be expressed in radians/second |
| absval\_rad\_s | same as absval\_rad, but multipied by 120 to be expressed in radians/second |
| dirval\_deg\_s | same as dirval\_deg, but multipied by 120 to be expressed in degrees/second |
| absval\_deg\_s | same as absval\_deg, but multipied by 120 to be expressed in degrees/second |
| dirval\_rad\_lr | same as dirval\_rad, but here positive values are for left, negative values are for right, to check for side bias |

### Analysis

Before proceeding with the analysis, a consideration. Due to hardware limitations, sometimes the rendering of stimuli slowed down, such that the stimulus speed would decrease. This means that the column “framefromonset” and “timefromonset” may be slightly disaligned in the table, as “framefromonset” is directly associated to the frame number, while “timefromonset” is taken from the computer timestamp.

```
ggplot(main, aes(x=framefromonset, timefromonset))+
geom_point(size=0.1)
```

From the graph one can clearly notice how this doesn’t happen often, and when it happens is generally for only one or two frames. Some stimuli have a more consistent delay, as one can see by the appearance of a second sloped line on the left of the graph. This happens because the script is constrained to produce frames once every 0.033 seconds, so if the software slows down it will produce a frame in the next available slot, effectively after a total of 0.066 seconds. This means that if a stimulus skips all frames, the slowest possible speed is 15fps. No further delays happens in the 30 seconds between two stimuli presentation, and in fact the points from then on run parallel to the expected line.

Crucially, the rotational position of the sphere are always correctly associated with the stimulus position for each given frame, independently from what time stamp information is used. The misalignment only cause the position of the stimulus to not match with different subjects according to the timefromonset column.

For this reason, from now on we will use the framefromonset column, such that there will be an exact correspondence for every individual between time and stimulus position.

#### Preliminary analysis

Before proceeding with the main analysis, we will perform some checks on the data

##### does stimulus position predicts absval?

Z peak values have been filtered by subtracting from it corresponding X and Y values. This has been done because we hypothesize that a rotation towards the left of the right stimulus would translate into a rotation of the treadmill sphere around its Z axis. This means that ideally during such a rotation, X and Y values should be 0. Due to noises of the system however, we cannot just delete Z peaks that have any X or Y concurrent values, as they are always present. We also cannot define a threshold, as it changes across subjects and across videos. By subtracting X and Y values, we purify the Z peak, so that now a higher peak is more likely to be an actual rotation rather than noise, and a lower peak should represent one that underwent more filtering. By doing this however we run the risk of losing the actual peak value: higher Z peaks should correspond to faster rotation, probably directed to further away stimuli. To check if our filtering did not denatured the real Z value, we will check if stimulus position predicts peak value. This will of course only be tested for when the stimulus is visible on the screen. For the analysis we will use the deg/sec versions of variables, as they are in a scale more immediately understandable. Note however that the scale should not change the results, as the transformation is linear and the variance is preserved in the change.

```
mori <- glmmTMB(absval_deg_s~stimpos_deg_all*cond + (1|subj/trialn),
                data = main, family = Gamma(link='log'),
                control=glmmTMBControl(optCtrl = list(iter.max = 30000, eval.max = 40000)))

simres <- simulateResiduals(mori)
plot(simres, factor=TRUE)
```

```
## Warning in plot.window(...): parametro grafico "factor" non valido
```

```
## Warning in plot.xy(xy, type, ...): parametro grafico "factor" non valido
```

```
## Warning in title(...): parametro grafico "factor" non valido
```

The deviation is significant, although the model seems fine. I have tried many other distributions, and this seems the best. Indeed, the full data is probably exponentially distributed (only positive, less and less values the higher they are), but here I am cutting only the during stimulus part. Will keep the model as is, it does not seem too critical here as it is only a check.

```
Anova(mori)
```

```
## Analysis of Deviance Table (Type II Wald chisquare tests)
## 
## Response: absval_deg_s
##                          Chisq Df Pr(>Chisq)    
## stimpos_deg_all      3330.8079  1  < 2.2e-16 ***
## cond                    2.5216  3     0.4714    
## stimpos_deg_all:cond   46.7756  3  3.879e-10 ***
## ---
## Signif. codes:  0 '***' 0.001 '**' 0.01 '*' 0.05 '.' 0.1 ' ' 1
```

Indeed, rotation depends on stimulus position. post-hoc follows

```
t <- emtrends(mori, ~cond, var='stimpos_deg_all')
pairs(t)
```

```
##  contrast                       estimate       SE    df t.ratio p.value
##  (bio-rand) - (bio-rigid)       0.000862 0.000795 20987  1.085  0.6990 
##  (bio-rand) - (rigid-rand)     -0.000276 0.000814 20987 -0.339  0.9866 
##  (bio-rand) - (shil-ellipse)   -0.004061 0.000793 20987 -5.123  <.0001 
##  (bio-rigid) - (rigid-rand)    -0.001138 0.000802 20987 -1.418  0.4883 
##  (bio-rigid) - (shil-ellipse)  -0.004923 0.000780 20987 -6.309  <.0001 
##  (rigid-rand) - (shil-ellipse) -0.003786 0.000800 20987 -4.730  <.0001 
## 
## P value adjustment: tukey method for comparing a family of 4 estimates
```

Correlation is positive: the more the stimulus is far away, the faster the rotation. Since saccadic rotations tend to generally have the same duration, a faster one means that the end point will be further away.

##### is dirval valuable?

To be sure that positive and negative values indeed suggest what the spider is turning towards the correct and the wrong stimulus, we designed a simple control condition: we presented a spider with a condition where only one stimulus at a time is present on the screen. for this condition dirval is positive for turns congruent with the position and negative for turns congruent with the empty side. We expect to observe an average peak value strongly positive, as the spider should only turn towards the side with the stimulus. We are not including any random effect, as it is only one subject. Indeed the p-value here is of marginal importance, the crucial point is that the results should be strongly positive.

```
mcontrol <- glmmTMB(dirval_deg_s~stimpres + (1|subj/trialn),
                    data = control, family = gaussian(link='identity'))
simres <- simulateResiduals(mcontrol)
plot(simres)
```

```
e <- emmeans(mcontrol, ~stimpres)
test(e, adjust='tukey')
```

```
##  stimpres emmean   SE  df t.ratio p.value
##  0         -24.5 13.9 119 -1.759  0.1557 
##  1         107.0 12.0 119  8.887  <.0001 
## 
## P value adjustment: sidak method for 2 tests
```

Indeed, when no stimulus is presented on the screen, the value is not different from zero, meaning there is no preference. When the stimulus is visible, the value is highly positive, signifying a higher number of rotation towards it.

Having now demonstrated that the peak magnitude is consistent with the stimulus position, and the fact that the sign of the peak is consistent with the stimulus side, we can proceed with the main analysis confident in the fact that we are actually measuring the turn preference.

#### Main Analysis

To test the preference of the spiders, we will include in the model:

- cond, to see the value in every condition.
- stimpres, to check the difference between when the stimulus is visible and when is absent. We expect that when the stimuli are not visible the average peak should be 0, as there should be an equal amount of rotation towards the left and the right (positive and negative)
- sex, to see if it has any effect on the conditions.
- stimulus number, to see if earlyer stimuli have something different to later ones.

```
mmain <- glmmTMB(dirval_deg_s~stimn*cond*stimpres + (1|subj/trialn),
                 data=main, family= gaussian(link = 'identity'),
                 control=glmmTMBControl(optCtrl = list(iter.max = 30000, eval.max = 40000)))

simres <- simulateResiduals(mmain)
plot(simres)
```

This distribution remains strange. Indeed, the reason is that it is actually representing an exponential distribution mirrored on the negative space to use the sign as an indication of correctness. We can’t analyze the two distribution separately and then test them against each other, because in that case we would only look at the difference of magnitudes, rather than overall preference: a spider turning once towards the correct stimulus and turning three times towards the wrong one, every time with the same magnitude, would show no difference as the average value of the wrongs would be the same as the average value of the correct. At the same time we cannot just use a count variable, as keeping the magnitude makes us able to give less weight to probable random noise and keeping high the weight of probable turns, without having to define a hard-coded threshold that would for certain cut some animals behaviour. For all these reasons, we will not try to fit the data into any given distribution, and test it as it is

```
Anova(mmain)
```

```
## Analysis of Deviance Table (Type II Wald chisquare tests)
## 
## Response: dirval_deg_s
##                       Chisq Df Pr(>Chisq)    
## stimn                0.5732  1  0.4489954    
## cond                 2.9164  3  0.4046989    
## stimpres            74.1006  1  < 2.2e-16 ***
## stimn:cond           2.4947  3  0.4762489    
## stimn:stimpres       1.8914  1  0.1690389    
## cond:stimpres       19.2137  3  0.0002469 ***
## stimn:cond:stimpres  2.4773  3  0.4793989    
## ---
## Signif. codes:  0 '***' 0.001 '**' 0.01 '*' 0.05 '.' 0.1 ' ' 1
```

```
e <- emmeans(mmain, ~cond*stimpres, type='response')
```

```
## NOTE: Results may be misleading due to involvement in interactions
```

```
e
```

```
##  cond         stimpres  emmean   SE    df lower.CL upper.CL
##  bio-rand     0          0.758 1.59 20979  -2.3539     3.87
##  bio-rigid    0          1.761 1.54 20979  -1.2522     4.78
##  rigid-rand   0          2.413 1.68 20979  -0.8784     5.70
##  shil-ellipse 0          3.445 1.73 20979   0.0628     6.83
##  bio-rand     1        -14.158 3.35 20979 -20.7152    -7.60
##  bio-rigid    1         -2.081 3.25 20979  -8.4456     4.28
##  rigid-rand   1        -15.685 3.36 20979 -22.2770    -9.09
##  shil-ellipse 1        -23.396 3.38 20979 -30.0229   -16.77
## 
## Confidence level used: 0.95
```

```
#i dont think this is needed
test(e, adjust='tukey')
```

```
##  cond         stimpres  emmean   SE    df t.ratio p.value
##  bio-rand     0          0.758 1.59 20979  0.477  0.9997 
##  bio-rigid    0          1.761 1.54 20979  1.146  0.9020 
##  rigid-rand   0          2.413 1.68 20979  1.437  0.7294 
##  shil-ellipse 0          3.445 1.73 20979  1.996  0.3133 
##  bio-rand     1        -14.158 3.35 20979 -4.232  0.0002 
##  bio-rigid    1         -2.081 3.25 20979 -0.641  0.9973 
##  rigid-rand   1        -15.685 3.36 20979 -4.664  <.0001 
##  shil-ellipse 1        -23.396 3.38 20979 -6.921  <.0001 
## 
## P value adjustment: sidak method for 8 tests
```

##### is there a side bias?

Finally, we need to test whether there is a pre-existing side bias. The position of the stimuli is counterbalanced across trials, so even if there is a side bias it will not influence the results of the experiment, but still it will be useful to know. Note that the distribution is the same, but will keep it as it is for the reasons described above

```
mside <- glmmTMB(dirval_rad_lr~stimpres*cond + (1|subj/trialn),
                 data = main, family = gaussian(link='identity'),
                control=glmmTMBControl(optCtrl = list(iter.max = 30000, eval.max = 40000)))

Anova(mside)
```

```
## Analysis of Deviance Table (Type II Wald chisquare tests)
## 
## Response: dirval_rad_lr
##                 Chisq Df Pr(>Chisq)  
## stimpres       4.0148  1    0.04510 *
## cond           2.2899  3    0.51447  
## stimpres:cond 10.2352  3    0.01667 *
## ---
## Signif. codes:  0 '***' 0.001 '**' 0.01 '*' 0.05 '.' 0.1 ' ' 1
```

```
e<-emmeans(mside, ~stimpres*cond, type='response')
e
```

```
##  stimpres cond            emmean       SE    df  lower.CL upper.CL
##  0        bio-rand      0.000463 0.000430 20987 -0.000379 0.001305
##  1        bio-rand      0.000467 0.000594 20987 -0.000696 0.001631
##  0        bio-rigid     0.000600 0.000414 20987 -0.000212 0.001412
##  1        bio-rigid    -0.000634 0.000577 20987 -0.001766 0.000498
##  0        rigid-rand   -0.000444 0.000445 20987 -0.001316 0.000428
##  1        rigid-rand    0.000158 0.000600 20987 -0.001018 0.001333
##  0        shil-ellipse  0.000499 0.000449 20987 -0.000382 0.001380
##  1        shil-ellipse -0.000982 0.000593 20987 -0.002144 0.000180
## 
## Confidence level used: 0.95
```

```
test(e, adjust='bonferroni')
```

```
##  stimpres cond            emmean       SE    df t.ratio p.value
##  0        bio-rand      0.000463 0.000430 20987  1.077  1.0000 
##  1        bio-rand      0.000467 0.000594 20987  0.787  1.0000 
##  0        bio-rigid     0.000600 0.000414 20987  1.448  1.0000 
##  1        bio-rigid    -0.000634 0.000577 20987 -1.098  1.0000 
##  0        rigid-rand   -0.000444 0.000445 20987 -0.999  1.0000 
##  1        rigid-rand    0.000158 0.000600 20987  0.263  1.0000 
##  0        shil-ellipse  0.000499 0.000449 20987  1.110  1.0000 
##  1        shil-ellipse -0.000982 0.000593 20987 -1.657  0.7804 
## 
## P value adjustment: bonferroni method for 8 tests
```

We see an effect of the stimulus presence and an effect of the interaction between it and the condition, but by applying a post-hoc we clearly see none of those is different from zero, presenting really small values, especially compared to the observed preference in the main analysis which was between +10 and +20. We can conclude there is no side bias.
