## Supplementary material for "Perception of biological motion in point-light displays by jumping spiders": Full analysis and python script: SI2-DataFilteringScript.py

```
"""
This script takes FicTrac output and transform it into the dataset analyzed in the paper.
This is done by:

    1. opening one single-trial csv
    2. applying Butterworth filter to all three axis (referenced to the spider point of view)
    3. Finding z-only (or almost z-only) movements, by subtracting absolute value of x and y to z,
       This way, we get a z line, the highest the value, the more the movement was purely on the Z axis.
       Z peaks that were lower than X or Y in the same time point get deleted.
    4. Finding peaks on this new purified Z
    5. Save all values in a new file

"""

import pandas as pd #to load and read csv
import os #to act on folders
import numpy as np #to do math operations
from scipy import signal #for filters

path = '/RawData/' #provide folder containing Raw data

tmp = os.listdir(path)
filelist = [file for file in tmp if file.endswith('FULL.csv')] #to be sure to get only the right data
del tmp


# prepare table where to save the results
results = []

#make header
results.append(['date', 'species', 'subj', 'sex', 'trialn', 'cond',
                'stimn', 'side', 'stimtype', 'turnamount', 'firstturn', 'choice'])

#main loop, do this for every file
for n, file in enumerate(filelist):
    print(str(n)+' - '+file)
    table = pd.read_csv(path+file)

    # get filter values and apply to x,y and z axis
    b, a = signal.butter(3, 0.05)
    table['DELTArvLabXfilt'] = signal.filtfilt(b, a, table['DELTArvLabX'].values)
    table['DELTArvLabYfilt'] = signal.filtfilt(b, a, table['DELTArvLabY'].values)
    table['DELTArvLabZfilt'] = signal.filtfilt(b, a, table['DELTArvLabZ'].values)

    #utilities to convert pixel positions to mm and then to angles
    table['leftstimposmm'] = table['leftstimpos'] * table['pixpitch']

    table['leftstimposmmExtremeLeft'] = (table['leftstimpos']-50) * table['pixpitch']
    table['leftstimposmmExtremeRight'] = (table['leftstimpos']+50) * table['pixpitch']

    table['leftstimposANGLE'] = np.arctan(table['leftstimposmm']/table['dist'])

    table['leftstimposANGLEExtremeLeft'] = np.arctan(table['leftstimposmmExtremeLeft'] / table['dist'])
    table['leftstimposANGLEExtremeRight'] = np.arctan(table['leftstimposmmExtremeRight'] / table['dist'])

    table['rightstimposmm'] = table['rightstimpos'] * table['pixpitch']

    table['rightstimposmmExtremeLeft'] = (table['rightstimpos']-50) * table['pixpitch']
    table['rightstimposmmExtremeRight'] = (table['rightstimpos']+50) * table['pixpitch']

    table['rightstimposANGLE'] = np.arctan(table['rightstimposmm']/table['dist'])

    table['rightstimposANGLEExtremeLeft'] = np.arctan(table['rightstimposmmExtremeLeft']/table['dist'])
    table['rightstimposANGLEExtremeRight'] = np.arctan(table['rightstimposmmExtremeRight']/table['dist'])

    table['time'] = table['timestamp_video']-table['timestamp_video'].values[0]

    #get binomial variable to tell if stimulus is present or not
    table['stimvisible'] = 0
    table['stimvisible'][table['framefromonset'].between(28,263)] = 1


    # routine to get Z axis with X and Y removed
    Zclean = []
    for n in range(len(table)) :
        z = table['DELTArvLabZfilt'].values[n]
        x = table['DELTArvLabXfilt'].values[n]
        y = table['DELTArvLabYfilt'].values[n]
        o = abs(z) - abs(x) - abs(y)

        if o < 0:
            o = 0 #if o got below 0, set it as 0
        if z < 0:
            o = o * -1 #this is done to re-apply sign to the absolute value
        Zclean.append(o)
    table['DELTArvLabZclean'] = Zclean
    table['DELTArvLabZAbsolute'] = abs(table['DELTArvLabZclean'])


    #now, to create the timefromonset column, specifying the time in term of distance from the stimulus appearance
    timefromonset = []
    starttimes = []
    for s in range(10): #once for every stimulus of the test
        currstim = table[table['stimn'] == s] #split the table and save the current stimulus
        starttimes.append(currstim['time'][currstim['stimvisible'] == 1].values[0]) #save the starting time

    # now for every row in the table
    for n in range(len(table)):
        if np.isnan(table['stimpres'].values[n]): #don't do anything if we are in the habit period
            timefromonset.append(np.nan)
        else:
            stimn = table['stimn'].values[n]
            currtime = table['time'].values[n] - starttimes[int(stimn)]
            timefromonset.append(currtime) #write the time as a function of stimulus onset
    table['timefromonset'] = timefromonset


    #find peaks on the filtered Z axis
    pos, val = signal.find_peaks(table['DELTArvLabZclean'], height=0.001)
    timepos = [table['time'].values[p] for p in pos]
    valpos = val['peak_heights']

    #also in the negative space
    pos, val = signal.find_peaks(table['DELTArvLabZclean'] * -1, height=0.001)
    timeneg = [table['time'].values[p] for p in pos]
    valneg = val['peak_heights'] * -1

    #save the timestamps of the peaks and the values, then put them in a dictionary together
    timeZ = np.concatenate((timepos, timeneg))
    valZ = np.concatenate((valpos, valneg))
    peaksZ = {'time': timeZ, 'val': valZ}
    peaksZ = pd.DataFrame(peaksZ)

    peaksZ = peaksZ.sort_values('time')
    timeZ = peaksZ.time.values
    valZ = peaksZ.val.values


    # #SAVING ROUTINE# #

    #get the data of the current trial from the filename
    date = table['date'].values[0]
    species = table['species'].values[0]
    subj = table['name'].values[0]
    sex = table['sex'].values[0]
    trialn = table['trialn'].values[0]

    #for every peak found (getting time and value)
    for time, val in zip(timeZ, valZ):
        first = 0
        oldstimn = -1
        if not table['stimn'][table['time'] == time].isnull().item():
            stimn = table['stimn'][table['time'] == time].item()

            #was the stimulus present during this peak?
            stimpres = table['stimvisible'][table['time'] == time].item()

            oldstimn = stimn
            leftstim = table['leftstimtype'][table['time'] == time].item()
            rightstim = table['rightstimtype'][table['time'] == time].item()
            stims = [leftstim, rightstim]
            #set the correct side depending on the two alternatives and also save the condition as factor
            if 'bio' in stims and 'rigid' in stims:
                cond = 'bio-rigid'
                if leftstim == 'bio':
                    corrside = 'left'
                else :
                    corrside = 'right'
            elif 'bio' in stims and 'rand' in stims:
                cond = 'bio-rand'
                if leftstim == 'bio':
                    corrside = 'left'
                else :
                    corrside = 'right'
            elif 'rand' in stims and 'rigid' in stims:
                cond = 'rigid-rand'
                if leftstim == 'rigid' :
                    corrside = 'left'
                else :
                    corrside = 'right'
            elif 'shil' in stims and 'ellipse' in stims:
                cond = 'shil-ellipse'
                if leftstim == 'shil' :
                    corrside = 'left'
                else :
                    corrside = 'right'

            elif 'ellipse' in stims:
                cond = 'control'
                if leftstim == 'ellipse':
                    corrside = 'left'
                else :
                    corrside = 'right'

            dirval_rad_lr = val
            if corrside == 'left':
                dirval_rad = val
            elif corrside == 'right':
                dirval_rad = val * -1
            if val > 0 :
                binom = 1
            elif val < 0 :
                binom = 0

            #change the variables scale: from radians to degrees, and multiply the delta for that frame by 120 to get peak speed in deg or rad per second
            absval_rad = abs(dirval_rad)
            dirval_deg = np.rad2deg(dirval_rad)
            absval_deg = np.rad2deg(absval_rad)
            dirval_rad_s = dirval_rad * 120
            absval_rad_s = absval_rad * 120
            dirval_deg_s = dirval_deg * 120
            absval_deg_s = absval_deg * 120

            if stimpres == 1:
                stimpos_rad = table['rightstimposANGLE'][table['time'] == time].item()
                stimpos_deg = np.rad2deg(table['rightstimposANGLE'][table['time'] == time].item())
            else:
                stimpos_rad = np.nan
                stimpos_deg = np.nan

            framefromonset = table['framefromonset'][table['time'] == time].item()
            timefromonset = table['timefromonset'][table['time'] == time].item()

            #add row
            results.append(
                [date, species, subj, sex, trialn, stimn, cond, corrside, stimpres, framefromonset, timefromonset,
                 stimpos_rad, stimpos_deg, dirval_rad, absval_rad, dirval_deg, absval_deg, dirval_rad_s, absval_rad_s,
                 dirval_deg_s, absval_deg_s, dirval_rad_lr])

#save
np.savetxt('results_FULL.csv',results, delimiter=",", fmt='%s')
```
